# Divergent immune trajectories distinguish resistance and susceptibility to bacterial leaf streak in wheat

**DOI:** 10.64898/2026.11.19.767916

**Authors:** Jatinder Singh, Santosh Gudi, Peter J. Maughan, Zhaohui Liu, Sudeshi Seneviratne, Katherine L.D. Running, Krishna Acharya, Andrew J. Green, Karl D. Glover, Upinder Gill, Justin D. Faris, Rajeev Gupta

## Abstract

Bacterial leaf streak (BLS) is an increasingly important disease affecting several cereal crops. The mechanisms linking quantitative trait loci for resistance to downstream immune response against BLS in wheat remain poorly understood. Here, we developed a chromosome-scale genome assembly of the BLS resistant cultivar Boost. With *de novo* annotation of Boost and Timstein (susceptible cultivar) assemblies, we established a cultivar-resolved framework for dissecting BLS resistance. Both cultivars encoded highly similar immune receptor repertoires, but the major resistance locus, *QBls.fcu-3B.1*, comprised a divergent receptor kinase haplotype with extensive structural variations, divergent tandem receptor arrays, and constitutively elevated expression of multiple receptor kinases in the resistant cultivar. Candidate receptor kinases in Boost exhibited conserved intracellular kinase domains but substantial divergence among extracellular recognition domains relative to their syntenic homologs in Timstein. Dual reference transcriptome analyses at 24, 48, and 96 hours after inoculation (HAI) revealing contrasting immune trajectories in resistant and susceptible cultivars. Boost progressed through a focused signaling transition at 48 HAI, followed by coordinated activation of several defense modules, such as antimicrobial metabolism, cell wall reinforcement, transcriptional regulation, and immune homeostasis. In contrast, the susceptible cultivar underwent coordinated repression of these defense-related modules together with chromatin-associated transcriptional changes and induction of host transcriptional machinery during disease establishment. Our findings suggest that quantitative resistance to BLS is associated not simply with immune receptor composition, but also with constitutive immune surveillance and temporal coordination of downstream defense, providing a putative mechanistic framework linking natural genomic variations to quantitative BLS resistance in wheat.

## Introduction

Plants rely on a multilayered innate immune system to detect and limit microbial invasions while maintaining their own growth and development (Huot et al., 2014). The immune response is initiated by pattern recognition receptors (PRRs) present on the cell surface. PRRs typically include receptor-like kinases (RLKs) and receptor-like proteins (RLPs), which perceive the conserved pathogen-or damage-associated molecular patterns (PAMPs/DAMPs) and activate pattern-triggered immunity (PTI) (Couto and Zipfel, 2016). When pathogens overcome PTI, intracellular immune receptors, including nucleotide-binding leucine-rich repeat (NLR) proteins subsequently recognize pathogen effectors to activate effector-triggered immunity (ETI) (Jones and Dangl, 2006). PTI and ETI reinforce defense responses through overlapping signaling networks (Ngou et al., 2021; Yuan et al., 2021). Although PTI and ETI are initiated by distinct receptor classes, both mechanisms converge on common downstream processes, including calcium influx, reactive oxygen species (ROS) production, mitogen-activated protein kinase (MAPK) signaling, transcriptional reprogramming, antimicrobial metabolite biosynthesis, cell wall reinforcement, and systemic acquired resistance (SAR) (Peng et al., 2018; Ramírez-Zavaleta et al., 2022; Yu et al., 2024). Consequently, successful disease resistance depends not only on pathogen recognition but also on precise temporal coordination of these downstream immune processes in the host (Li et al., 2020).

Immune receptor genes are among the most dynamic components in the plant genome. RLKs, RLPs, and NLRs frequently occur in tandem arrays that undergo duplication and recombination, resulting in copy-number variation and rapid sequence diversification, which generate highly polymorphic genomic regions serving as reservoirs of disease resistance diversity (Michelmore and Meyers, 1998; Barragan and Weigel, 2021; Sutherland et al., 2025). These rapidly evolving and structurally variable receptor clusters often underlie quantitative resistance to diverse pathogens. However, the identification of the causal genes and understanding the mechanisms governing immune signaling remain major challenges, especially for polyploid crops (Wersch and Li, 2019; Dievart et al., 2020; Li et al., 2025). Structural polymorphisms, divergent alleles, and complex regulatory networks often obscure the relationship between genotype and phenotype (Krattinger and Keller, 2016). Recent data suggest that quantitative disease resistance is rarely explained by the presence or absence of individual immune receptors. Instead, receptor diversification at structurally variable loci, combined with regulatory variation that shapes downstream immune deployment, appears to determine the outcome of host-pathogen interactions (Corwin and Kliebenstein, 2017). However, the genomic basis linking receptor diversity to downstream immune signaling and transcriptional reprogramming remains poorly understood (Wulff and Krattinger, 2022; Ngou et al., 2022).

Bacterial leaf streak (BLS), caused by *Xanthomonas translucens* pv. *undulosa* (Xtu), has become an increasingly important bacterial disease of wheat and barley in many cereal producing areas worldwide (Sapkota et al., 2020). Following entry through natural openings or wounds, the bacteria colonize the leaf apoplast, producing characteristic water-soaked lesions that expand into translucent streaks and ultimately necrotic tissue, reducing the photosynthetic capacity and thus grain yield (Friskop et al., 2023; Heiden et al., 2023). Like other *Xanthomonas* species, Xtu employs a type III secretion system to deliver effector proteins into the host cells, including transcription activator-like (TAL) effectors that manipulate host gene expression to promote bacterial colonization (Peng et al., 2019). Despite the agricultural importance of BLS, the molecular mechanisms underlying quantitative BLS resistance in wheat remain largely unknown (Sapkota et al., 2020). Genetic analyses have revealed quantitative control of BLS resistance in wheat, with resistance largely determined by additive-effect quantitative trait loci (QTL) (Acharya et al., 2024). In that study, two QTL, *QBls.fcu-3B.1* and *QBls.fcu-5A* for BLS resistance were identified with the former being a major QTL under controlled environment conditions (Acharya et al., 2024). However, translating these QTL into candidate genes has been hindered in part by the complexity of the hexaploid wheat genome.

Recent advances in long-read sequencing have enabled the generation of chromosome-scale assemblies that capture structural variations at immune loci with unprecedented resolution (Athiyannan et al., 2022). When integrated with comparative genomics and time-resolved transcriptomics, these resources provide an opportunity to move beyond QTL mapping and identify candidate resistance mechanisms underlying quantitative disease resistance. Here, we generated a chromosome-scale genome assembly and evidence-based annotation for the BLS-resistant cultivar Boost and reannotated the published genome assembly of the BLS-susceptible cultivar Timstein (Singh et al., 2026) using the same pipeline to enable direct comparative genomic and BLS-responsive transcriptomic analyses. By integrating comparative genomics, resistance gene analogs (RGAs) profiling, and BLS regulated RNA sequencing at 24, 48, and 96 hours after inoculation (HAI), we reconstructed the distinct immune trajectories underlying resistance and susceptibility for BLS. This study bridges the gap between quantitative resistance loci and their underlying molecular mechanisms and provides a putative framework for understanding how coordinated immune regulation shapes quantitative BLS resistance in wheat.

## Results

### Chromosome-level genome assemblies establish a cultivar-resolved comparative framework to dissect BLS resistance

Boost has been previously reported as BLS resistant cultivar which carry major BLS resistance loci on chromosome 3B (*QBls.fcu-3B.1*) and chromosome 5A (*QBls.fcu-5A*) (Acharya et al., 2024). To investigate the genomic basis of BLS resistance in Boost, we developed a chromosome-scale genome assembly using PacBio HiFi and Hi-C sequencing and compared it with the genome of the susceptible cultivar Timstein (Singh et al., 2026) (Figure 1A, B, Table S1). The primary contig assembly of Boost was scaffolded into 21 chromosome-level pseudomolecules with total genome size of 14.58 Gb (Table S2 and S3; Figure S1; Note S1). Assembly quality was high with sub-genome BUSCO completeness ranging from 97.7-99.6%, LTR assembly index (LAI) of 13.7-18.1, k-mer completeness of 98.8% and a consensus quality value (QV) of 67.4 (Table S3). A total of 98.31% of the assembly was anchored to 21 chromosomes, with 3B representing the largest chromosome (863.76 Mb), consistent with other bread wheat genomes (Table S4). Merqury QV of 73.9-81.9 and recovery of 31 of 42 telomeric chromosome ends demonstrated high base-level accuracy and near-complete chromosomal representation (Table S5 and S6). As expected for wheat genomes, transposable elements (TEs) occupied 85.82% of the total Boost genome (Table S7).

**Figure 1.**
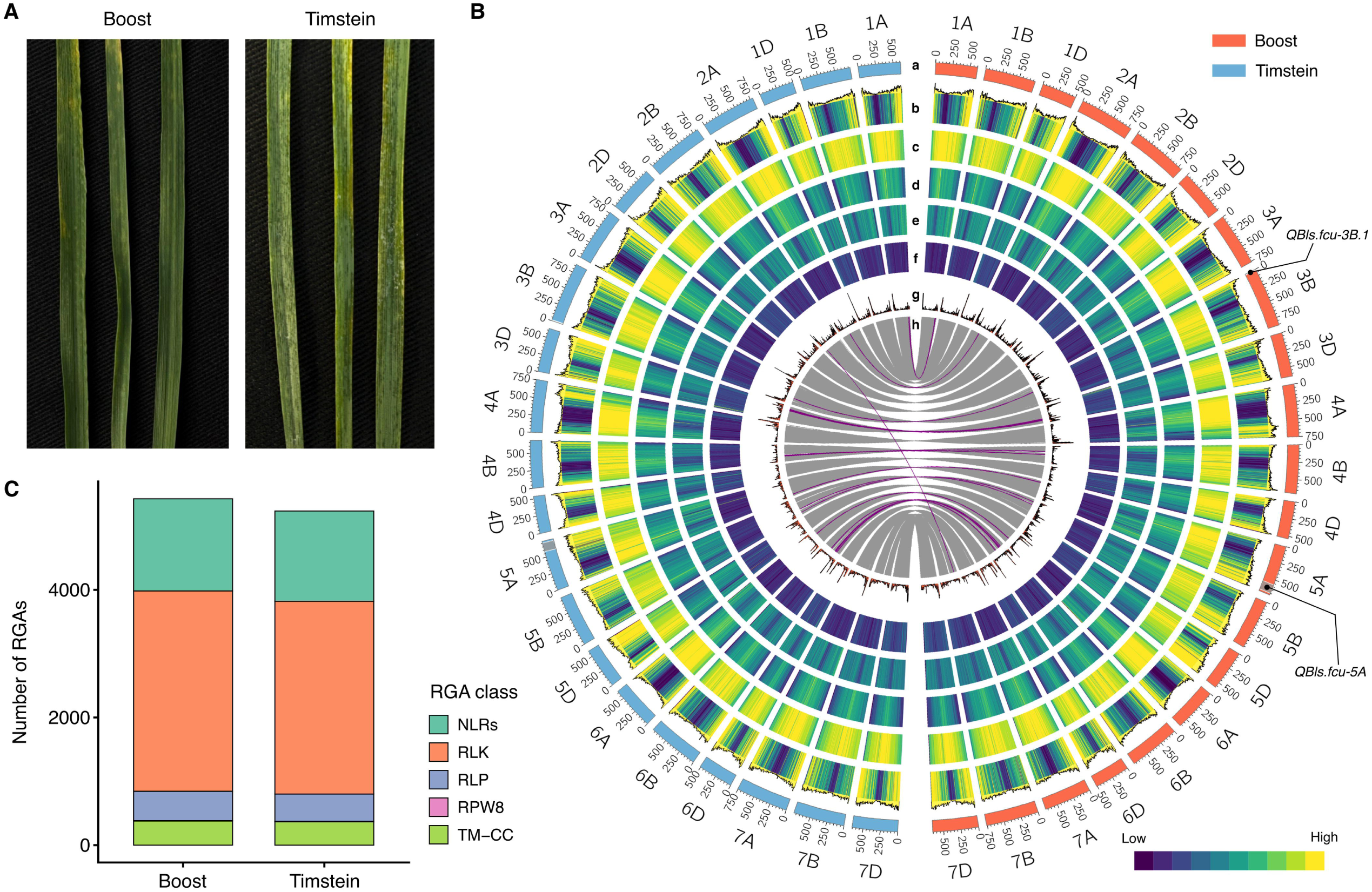
Chromosome-scale comparative genomics of the bacterial leaf streak-resistant (Boost) and susceptible cultivar (Timstein). **(A)** Representative leaves of Boost (left panel) and Timstein (right panel) following inoculation with *Xanthomonas translucens* pv. *undulosa* (Xtu), showing bacterial leaf streak symptoms in Timstein relative to Boost. **(B)** Circos representation of the chromosome-scale Boost and Timstein genome assemblies. From the outermost to the innermost tracks: (a) chromosome ideograms and physical coordinates in megabases for Boost and Timstein; (b) high-confidence gene density, displayed as a heatmap with an accompanying histogram; (c) repeat density; (d) Gypsy long terminal repeat retrotransposon density; (e) Copia long terminal repeat retrotransposon density; (f) CACTA DNA transposon density; (g) resistance gene analogue (RGA) density; and (h) syntenic relationships between high-confidence genes in Boost and Timstein. Grey ribbons represent collinear relationships, whereas purple ribbons indicate inverted relationships. The positions of the previously identified BLS resistance loci *QBls.fcu-3B.*1 and *QBls.fcu-5A* are indicated on the Boost chromosomes. **(C)** Comparison of high-confidence resistance gene analogs (RGAs) repertoires in Boost and Timstein. Stacked bars show the numbers of receptor-like kinases (RLKs), nucleotide-binding leucine-rich repeat receptors (NLRs), receptor-like proteins (RLPs), RPW8-domain proteins, and transmembrane coiled-coil proteins (TM–CCs).

Boost (resistant to BLS) and Timstein (susceptible to BLS) genome (Singh et al., 2026) assemblies were annotated using the same evidence guided BRAKER3 workflow, ensuring that downstream differences reflect biological variations rather than annotation methodology. We found 116,881 and 117,404 high-confidence (HC) gene models in Boost and Timstein, respectively (Table S8). BUSCO analysis of the predicted protein-coding genes demonstrated high annotation completeness, with both cultivars recovering more than 99% of conserved Poales orthologs (Figure S2). Gene structure characteristics, including exon number, coding sequence length, and monoexonic genes, were highly similar between Boost and Timstein (Table S8). Genome-wide resistance gene analog (RGA) annotations identified 5,427 and 5,236 RGAs in Boost and Timstein, respectively. Boost was found to have modestly more RLKs and RLPs than Timstein, but the overall immune receptor repertoires were highly similar between the two cultivars (Figure 1C; Table S9 and S10).

Whole genome alignment of the Boost genome assembly exhibited structural variants (SVs) despite overall chromosome collinearity with Timstein (Singh et al., 2026) and CS-IAAS (Liu et al., 2025) genomes (Figure 1B and Figure S3A-C). Consistent with their shared North American origin, Boost showed substantially fewer sequence and structural differences relative to Timstein than to the Chinese Spring (CS-IAAS) reference (Table S11). Compared with CS-IAAS, Boost contained approximately one-third more sequence and structural variants than in the Boost-Timstein comparison, including more SNPs (20.5 vs. 15.0 million), insertions, deletions, inversions, and translocations, indicating a considerably closer genomic relatedness between the two cultivars. Further, we found that the inversions were not evenly distributed across the three sub-genomes. In both comparisons, i.e., Boost-CS and Boost-Timstein, the B sub-genome contained the largest proportion of inversions, accounting for 54% and 58% of the observed inversions, respectively. Whereas the D sub-genome contained only 6% and 10% of the inversions, respectively (Table S12). Collectively, these chromosome-level genome assemblies establish a cultivar-resolved comparative genomic framework in wheat.

### A structurally and transcriptionally distinct receptor kinase haplotype is associated with QTL *QBls.fcu-3B.1*

The previously reported major QTL, *QBls.fcu-3B.1*, spanned approximately 8.47 Mb in Boost (6.35-14.83 Mb) and 8.11 Mb in Timstein (6.38-14.49 Mb) (Figure 2A; Table S13). In the Boost assembly, the QTL interval contained 214 high confidence (HC) gene models, whereas the syntenic homologous interval in the Timstein assembly had 160 HC genes, despite their comparable physical sizes (Table S14 and S15). Comparative analysis of the Boost and Timstein annotations revealed pronounced differences in the organization of the two haplotypes. Although the region was largely collinear, it varied in local gene organization, tandem receptor arrays, and structural variation. Structural variant (SV) analysis identified several SVs such as insertions, deletions, duplications, inversions, and translocations in this interval (Note S2), suggesting that *QBls.fcu-3B.*1 may represent a structurally divergent genomic haplotype between these genotypes. Inspection of annotated genes revealed that the two haplotypes differ in the composition and arrangement of multiple tandem gene arrays, including receptor-like kinases (RLKs), resistance proteins, germin-like proteins, expansins, and several families encoding proteins with unknown function. Rather than exhibiting the simple expansion or contraction of a single immune receptor family, the QTL region comprises structurally unique genomic haplotypes with different local gene architectures.

**Figure 2.**
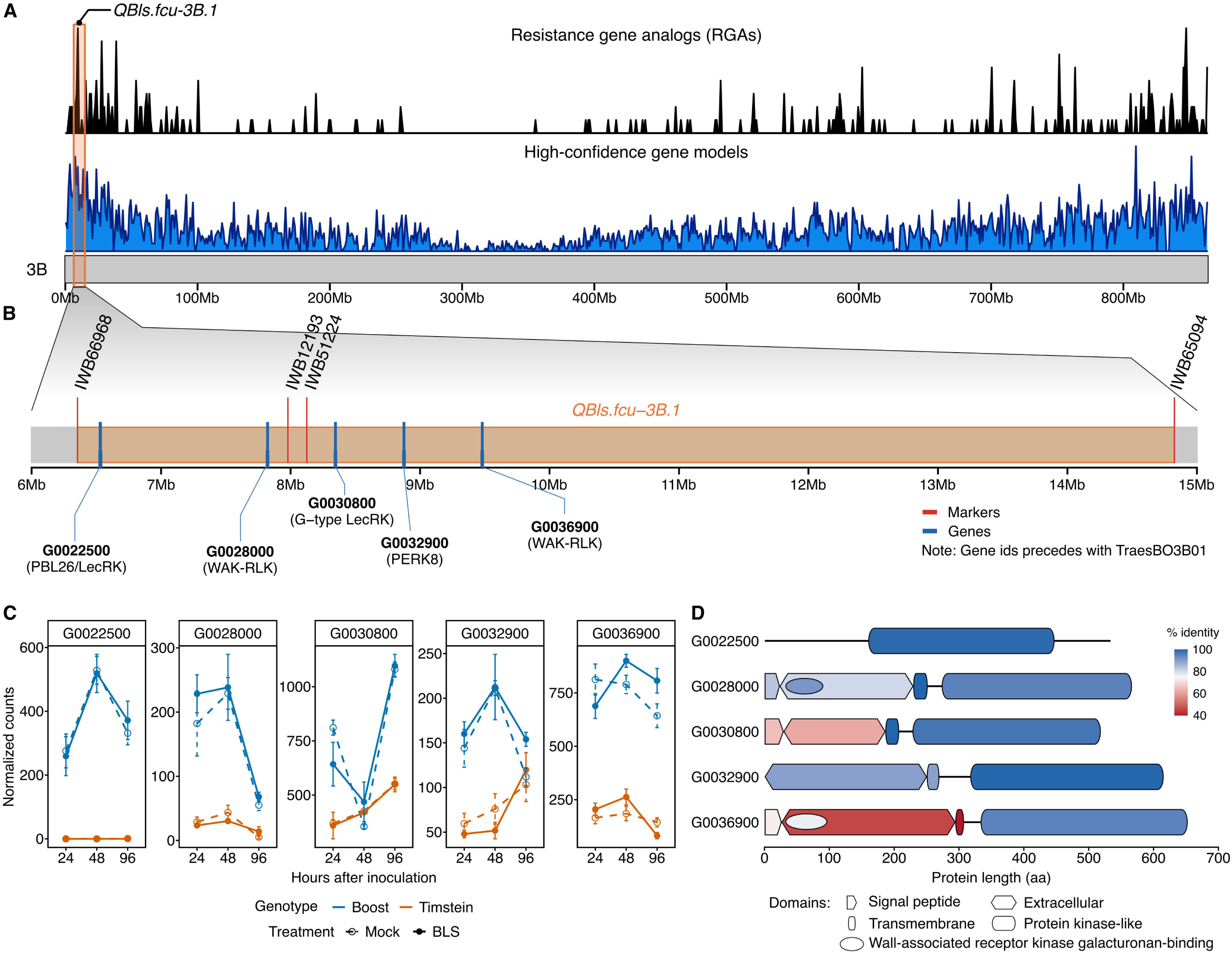
The major bacterial leaf streak resistance locus *QBls.fcu-3B.1* contains a receptor kinase-rich candidate interval. **(A)** Distribution of resistance gene analogs (RGAs) and high-confidence gene models along chromosome 3B of Boost. The physical position of *QBls.fcu-3B.1* is highlighted near the chromosome terminus. **(B)** Physical organization of the approximately 6–15-Mb *QBls.fcu-3B.1* interval. Red vertical lines indicate genetic markers, and blue vertical lines indicate annotated genes. The interval is bound by markers *IWB66968* and *IWB65094*. Candidate gene identifiers are abbreviated by removing the *TraesBO3B01* prefix; predicted functions are shown in parentheses. **(C)** Expression profiles of five prioritized Boost candidate genes and their Timstein orthologues under mock and BLS-inoculated conditions at 24, 48, and 96 hours after inoculation (HAI). Points represent mean DESeq2-normalized counts, and error bars represent the standard error of three biological replicates. Colors indicate cultivar, and solid and dashed lines distinguish BLS-inoculated and mock-treated samples, respectively. **(D)** Predicted protein-domain architectures of the five candidates in Boost. Domain colors represent amino acid identity relative to the corresponding Timstein ortholog. Predicted features include signal peptides, extracellular regions, transmembrane helices, protein kinase-like domains, and wall-associated receptor kinase galacturonan-binding domains. Protein coordinates are shown in amino acids.

No genes within the *QBls.fcu-3B.1* interval were differentially expressed in response to BLS infection in Boost. Therefore, we compared differential gene expression between BLS inoculated Boost and Timstein at 24, 48, and 96 hours after inoculation (HAI) (Table S16). In the *QBls.fcu-3B.1* interval, 22 HC genes were differentially expressed (16 up-and 6 down-regulated) between the two cultivars at one or more time points (Table S17). Among the 22 differentially expressed genes (DEGs), receptor-like kinases represented the largest defense-related gene class. Five receptor kinase genes were prioritized as candidate genes because they consistently exhibited higher transcript abundance in Boost across mock and inoculated conditions, encoded intact receptor proteins, and localized within the structurally divergent receptor-rich haplotype (Figure 2B). Five receptor kinase candidates consistently showed higher normalized expression counts in Boost (Figure 2C). We observed two wall-associated receptor kinases (WAK), *TraesBO3B01G0036900* and *TraesBO3B01G0028000*, maintaining consistently higher transcript abundance in Boost than Timstein. Additional receptor kinase candidates, such as a G-type lectin receptor kinase (*TraesBO3B01G0030800*), a proline-rich receptor-like protein kinase (*TraesBO3B01G0032900*), and a serine/threonine-protein kinase (*TraesBO3B01G0022500*) also exhibited elevated expression in Boost (Figure 2C). Further, protein sequence comparisons demonstrated that these transcriptional differences were accompanied by selective structural divergence. We observed that intracellular kinase domains remained highly conserved (94.7-99.7% amino acid identity), whereas extracellular domains showed substantially higher sequence divergence (45.5-89.6% amino acid identity) (Figure 2D; Table S18). This pattern was particularly evident for a WAK-RLK, *TraesBO3B01G0036900*, in which the conserved intracellular signaling domain contrasted with highly divergent extracellular regions likely involved in ligand perception. Such domain specific diversification is consistent with evolutionary modification of pathogen-recognition interfaces while maintaining downstream signaling capacity.

We also characterized the expression profile of genes within the BLS resistance locus, *QBls.fcu-5A*, on chromosome 5A. The QTL region spanned from 568.46 Mb to 698.28 Mb in Boost. At 24 and 48 HAI, no gene was differentially expressed between BLS-inoculated and mock-inoculated Boost samples. At 96 HAI, Boost showed substantial transcriptional activation, where a total of seven DEGs were identified, of which six were upregulated (Table S19 and S20). The upregulated genes encode defense-related proteins involved in membrane trafficking, calcium signaling, antimicrobial activity, and cell wall reinforcement. While the downregulated gene, *TraesBO5A01G0720200*, encodes a NAC transcription factor. This locus showed significant induction of genes encoding pathogenesis-related protein, calmodulin-binding proteins, receptor-like kinases, and a DIMBOA UDP-glucosyltransferase, suggesting the coordinated activation of multiple defense related pathways at the later stage of infection.

The combination of localized structural diversification, constitutively elevated receptor kinase expression, and extracellular domain divergence suggests that *QBls.fcu-3B.1* may function primarily at the level of pathogen perception, whereas *QBls.fcu-5A* may contribute towards downstream defense responses. We further investigated the genome-wide immune programs activated following BLS infection to determine downstream defense responses.

### Boost progresses through a sequential immune program during BLS infection

We compared the transcriptional response of BLS-inoculated and mock-inoculated Boost plants at 24, 48, and 96 HAI (Table S16 and S19; Figure S4). The differential expression analysis identified 54, 12, and 314 DEGs at 24, 48, and 96 HAI, respectively (Figure 3A; Table S20). By 96 HAI, approximately threefold more induced genes were observed compared to repressed genes. DEGs were broadly distributed across the A, B, and D sub-genomes (Figure 3B). This pattern indicates that the resistant response in Boost was associated with late, coordinated genome-wide transcriptional reprogramming. We assigned the DEGs to seven defense-related modules representing successive stages of plant immunity (Figure 3C; Table S20 and S21). These biological modules revealed a temporal progression from early immune perception to defense activation and subsequent immune regulation (Figures 3D-J).

**Figure 3:**
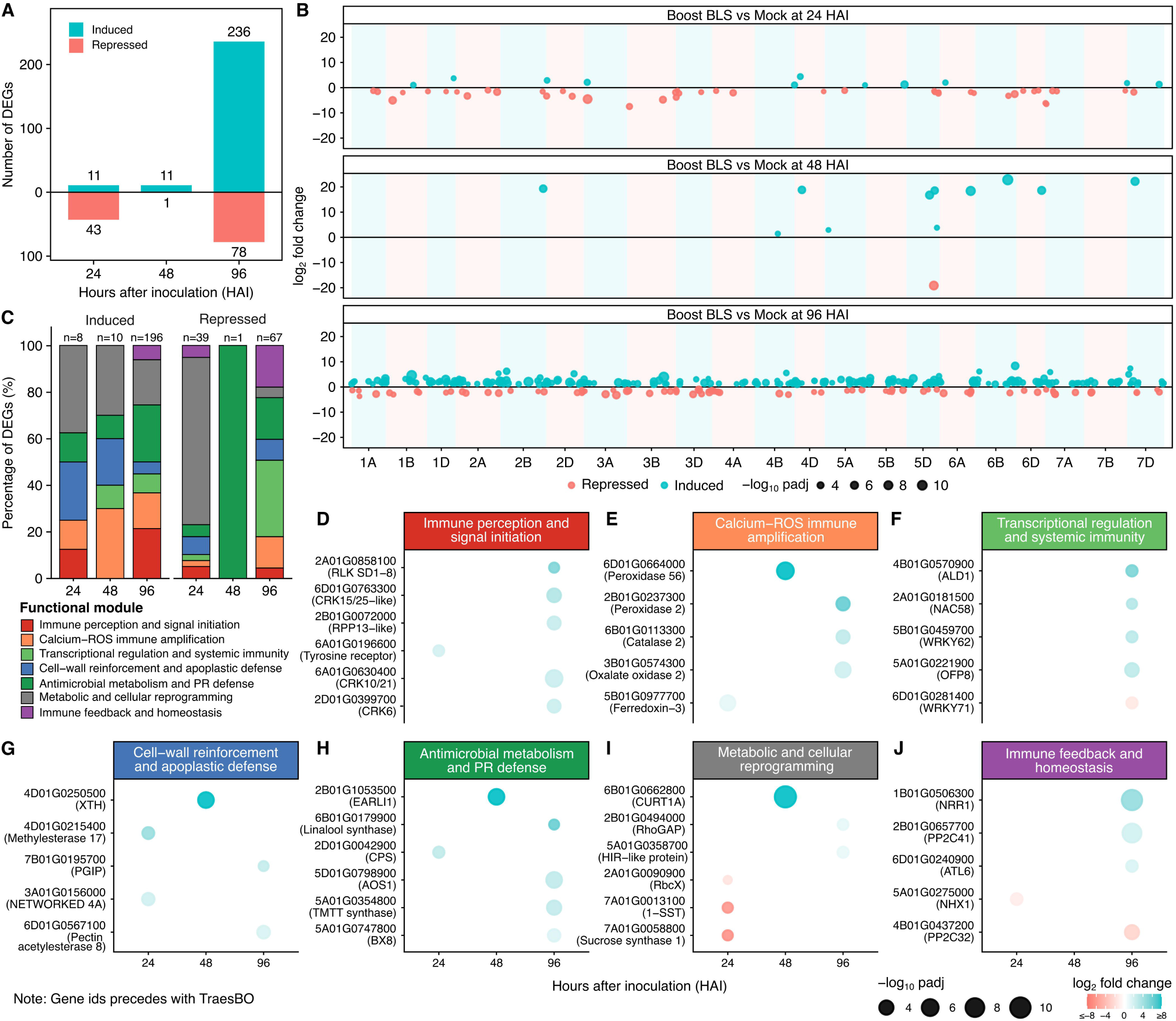
Temporal organization of the immune response in the bacterial leaf streak-resistant cultivar Boost. **(A)** Numbers of induced and repressed differentially expressed genes (DEGs) in Boost at 24, 48, and 96 hours after inoculation (HAI) with *Xanthomonas translucens* pv. *undulosa* relative to mock-treated controls. **(B)** Genome-wide distribution of DEGs across the 21 Boost chromosomes at each time point. Each point represents one DEG positioned according to its chromosomal location. Point size represents statistical significance (-log_10_ adjusted *P* value), and color indicates induction or repression. **(C)** Proportional distribution of induced and repressed DEGs among seven defense-associated functional modules and the Other/unknown category. Numbers above the bars indicate the total number of classified DEGs at each time point and regulatory direction. Temporal expression of representative genes associated with **(D)** immune perception and signal initiation, **(E)** calcium–ROS immune amplification, **(F)** transcriptional regulation and systemic immunity, **(G)** cell-wall reinforcement and apoplastic defense, **(H)** antimicrobial metabolism and pathogenesis-related defense, **(I)** metabolic and cellular reprogramming, and **(J)** immune feedback and homeostasis. Bubble size represents-log_10_ adjusted *P* value, and color represents log_2_ fold change. Gene identifiers are abbreviated by removing the *TraesBO* prefix.

At 24 HAI, the transcriptional response was limited in magnitude and was dominated by repression of primary metabolic and cellular processes. This early reduction in metabolic activity occurred alongside induction of a small set of receptor-like kinases, cell wall reinforcement, antimicrobial, and signaling components. The coexistence of restrained metabolism and selective defense activation indicates that Boost initiated immune perception before extensive downstream defense reprogramming. At 48 HAI there were only 12 DEGs, but they were qualitatively distinct from the response at 24 HAI. This stage represented a transition from pathogen perception to immune signal amplification. We identified DEGs enriched for ROS-associated signaling, immune transcriptional regulation, lipid transfer activity, and early cell wall modification (Table S20). Strong induction of a peroxidase and a germin-like protein indicated activation of oxidative processes (Figure 3E), while the induction of a xyloglucan endotransglycosylase/hydrolase (*XTH*) indicated cell wall modification (Figure 3G).

All seven defense-related modules were represented among the induced genes at 96 HAI. The immune perception module included several RLKs, cysteine-rich receptor kinases (CRKs), lectin receptor kinases (LRKs), and WAKs (Figure 3D; Table S20). This continued receptor activation was accompanied by broad reprogramming of calcium and ROS signaling, including induction of calmodulin-binding regulators, oxalate oxidases, and peroxidases (Figure 3E; Table S20 and S22). In contrast, several calcium-binding proteins were repressed, suggesting selective reconfiguration of calcium signaling rather than uniform activation of the entire gene family. Signal amplification was followed by extensive transcriptional reprogramming. Multiple WRKY, NAC (NAM, ATAF1/2, and CUC2), MYB (myeloblastosis), and bHLH (basic helix–loop–helix) transcription factor (TF) families were induced together with *ALD1* (*AGD2-LIKE DEFENSE RESPONSE PROTEIN 1*), suggesting activation of regulatory networks associated with both local and systemic defense (Figure 3F). Downstream responses expanded in parallel across complementary defense processes. Cell wall reinforcement involved polygalacturonase inhibitors, xyloglucan-modifying enzymes, methylesterases, pectin-modifying proteins and *WIR1*, consistent with strengthening of the physical barrier (Figure 3G). Antimicrobial defense involved coordinated activation of terpenoid, jasmonate, and benzoxazinoid metabolism, together with pathogenesis-related (PR) and detoxification proteins (Figure 3H). Defense activation was also accompanied by substantial cellular and metabolic reprogramming (Figure 3I; Table S20 and S22). Genes associated with immune regulation and homeostasis, such as *NRR1*, *ATL6*, and *PP2C41*, were induced, whereas three homoeologous copies of *PP2C32* were repressed (Figure 3J). The contrasting regulation of phosphatase subfamilies was consistent with dynamic control of phosphorylation-dependent defense signaling and immune homeostasis. Several of the most strongly induced genes encoded proteins of unknown function, indicating that additional components of BLS response remain uncharacterized (Table S20).

These results suggest that Boost progresses through a temporally coordinated immune programming during BLS infection that may lead to a resistant response. This biological progression from pathogen perception to defense deployment provides a framework for comparison with the susceptible response.

### Timstein undergoes immune suppression and transcriptional dysregulation during disease progression

To characterize transcriptional processes associated with susceptibility, we compared BLS-inoculated and mock Timstein plants at the same time points examined in Boost (Table S23). Differential expression analysis identified 102, 265, and 106 DEGs at 24, 48, and 96 HAI, respectively (Figure 4A; Table S24). In contrast to the late induction-dominated response in Boost, the largest Timstein response occurred at 48 HAI, which was dominated by repression of 199 DEGs and only 66 genes were induced. Across the three time points, 29 of 33 DEGs assigned to the immune perception and signal initiation modules were repressed. Susceptibility was associated not with transcriptional unresponsiveness, but with extensive negative regulation during disease establishment. The DEGs were distributed across A, B, and D sub-genomes (Figure 4B). Using the seven defense-related modules, we observed that Timstein exhibited a distinct temporal trajectory characterized by early alterations in immune signaling, widespread suppression during BLS infection, and a comparatively limited late response (Figure 4C-4J; Table S24).

**Figure 4:**
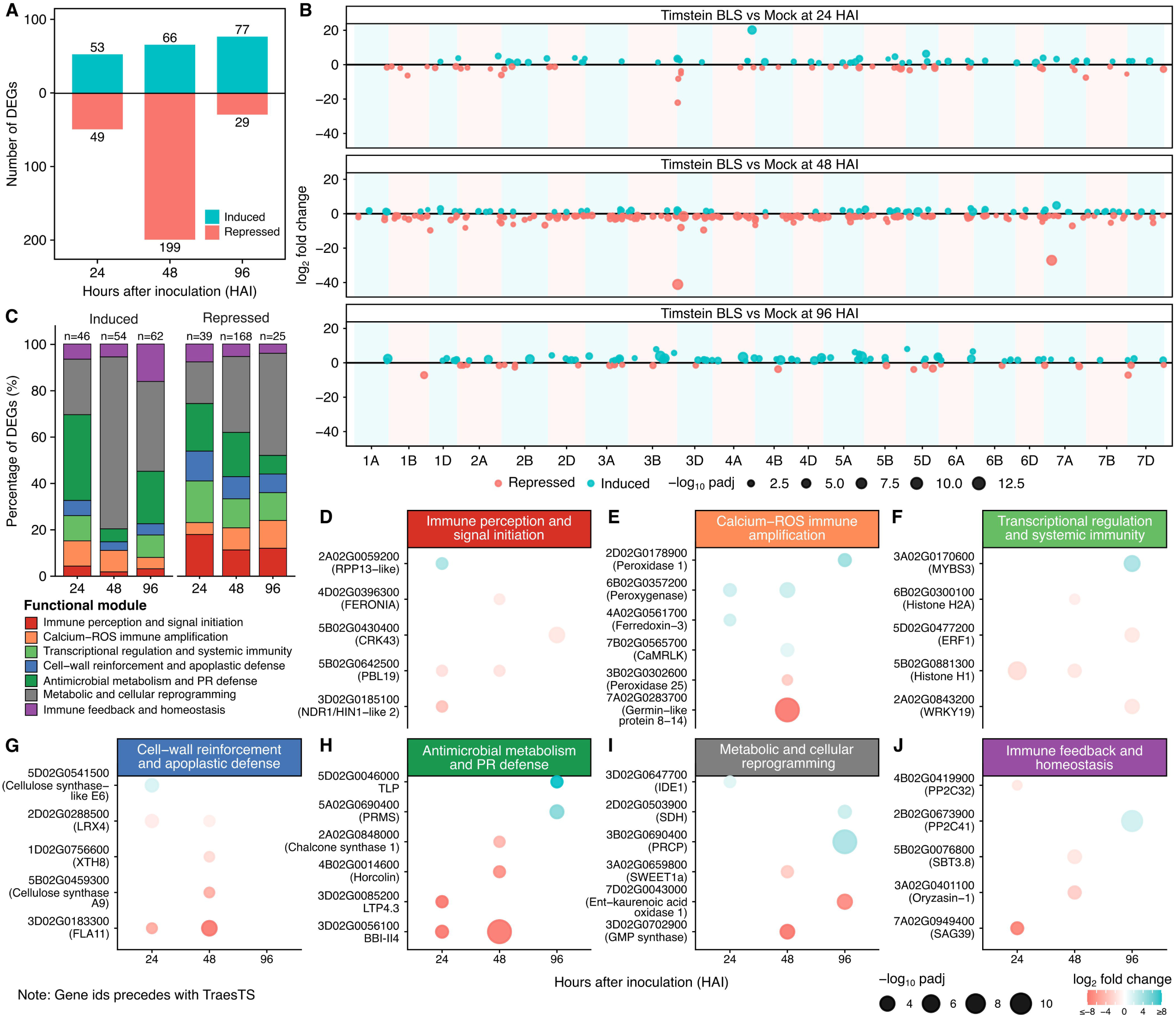
Temporal transcriptional dysregulation in the bacterial leaf streak-susceptible cultivar Timstein. **(A)** Numbers of induced and repressed differentially expressed genes (DEGs) in Timstein at 24, 48, and 96 hours after inoculation (HAI) with *Xanthomonas translucens* pv. *undulosa* relative to mock-treated controls. **(B)** Genome-wide distribution of DEGs across the 21 Timstein chromosomes at each time point. Each point represents one DEG positioned according to its chromosomal location. Point size represents statistical significance (-log_10_ adjusted *P* value), and color indicates induction or repression. **(C)** Proportional distribution of induced and repressed DEGs among seven defense-associated functional modules and the Other/unknown category. Numbers above the bars indicate the total number of classified DEGs at each time point and regulatory direction. Temporal expression of representative genes associated with **(D)** immune perception and signal initiation, **(E)** calcium–ROS immune amplification, **(F)** transcriptional regulation and systemic immunity, **(G)** cell-wall reinforcement and apoplastic defense, **(H)** antimicrobial metabolism and pathogenesis-related defense, **(I)** metabolic and cellular reprogramming, and **(J)** immune feedback and homeostasis. Bubble size represents - log_10_ adjusted *P* value, and color represents log_2_ fold change. Gene identifiers are abbreviated by removing the *TraesTS* prefix.

At 24 HAI, induced and repressed genes were present in similar numbers (Figure 4A), but several receptor and signaling components were already repressed, including *NDR1/HIN1*-like (NON-RACE-SPECIFIC DISEASE RESISTANCE 1 / HARPIN-INDUCED 1-like) and protein kinase *PBL19* (Figure 4D). The early response also involved coordinated repression of chromatin-associated components. Seven linker histone H1 genes were downregulated at 24 HAI, suggesting that altered chromatin regulation or nucleosome assembly occurred early in the susceptible interaction (Figure 4F; Table S24 and S25). Genes involved in antimicrobial activity and cell wall reinforcement were also strongly repressed during the early stages of BLS infection (Table S24 and S26).

The most pronounced transcriptional changes occurred at 48 HAI, when repression expanded across all seven defense-related modules. Genes involved in immune perception, calcium-ROS signaling, transcriptional regulation, antimicrobial metabolism, cell wall reinforcement, defense-associated cellular reprogramming and immune homeostasis were simultaneously repressed (Figure 4D-4J; Table S24). In contrast, genes associated with ribosome biogenesis, RNA processing, and translation were induced, indicating substantial reorganization of host transcriptional activity during disease establishment (Table S24 and S25). At 96 HAI, transcriptional response remained comparatively limited and was characterized by selective rather than coordinated defense activation. Several defense-associated genes, including thaumatin-like protein (TLP), endochitinase, pathogenesis-related protein, peroxidase, and CRKs, were strongly induced. However, these responses occurred alongside continued repression of other immune receptors and signaling components, including WAK2 and LRR-RLKs, indicating that immune activation remained fragmented rather than integrated (Table S24).

These results suggest that Timstein does not simply exhibit a weaker form of Boost response. Instead, the susceptible cultivar follows a distinct transcriptional program characterized by early attenuation of receptor and defense-related processes during disease establishment and a fragmented late defense response.

### Divergent immune trajectories distinguish resistant and susceptible cultivars

In direct comparison of the resistant and susceptible responses, we examined the temporal organization of the seven defense-related functional modules across both cultivars (Figure 5A). Both cultivars responded transcriptionally to Xtu infection, but the timing and organization of defense-associated processes varied substantially. Boost showed progressive activation of defense-related modules over time. On the contrary, Timstein showed extensive repression of these same modules during the middle stages of infection (Figure 5A). The greatest divergence between cultivars occurred during the 48 HAI transition. Only a small number of genes were differentially expressed in Boost, but those were highly enriched for calcium signaling, ROS generation, immune transcriptional regulation, and early cell wall modification (Figure 5B). This represents a highly focused signaling phase, which is followed by the broad defense activation observed at 96 HAI. In the case of Timstein, coordinated repression across all defense-related processes occurred at 48 HAI, which was associated with the susceptible response (Figure 5B). This repression occurred concurrently with increased expression of genes involved in ribosome biogenesis, RNA processing, and translation, suggesting a major reorganization of host transcriptional activity. The consequences of these contrasting intermediate responses became apparent at 96 HAI. Boost showed coordinated activation across all defense-related processes. Timstein also induced several defense-related genes, but these responses remained limited and fragmented across functional modules (Figure 5A and 5B). This contrasting organization was particularly evident in the antimicrobial activity and secondary metabolism, pathogen perception and immune receptors, and cell wall reinforcement sub-modules.

**Figure 5:**
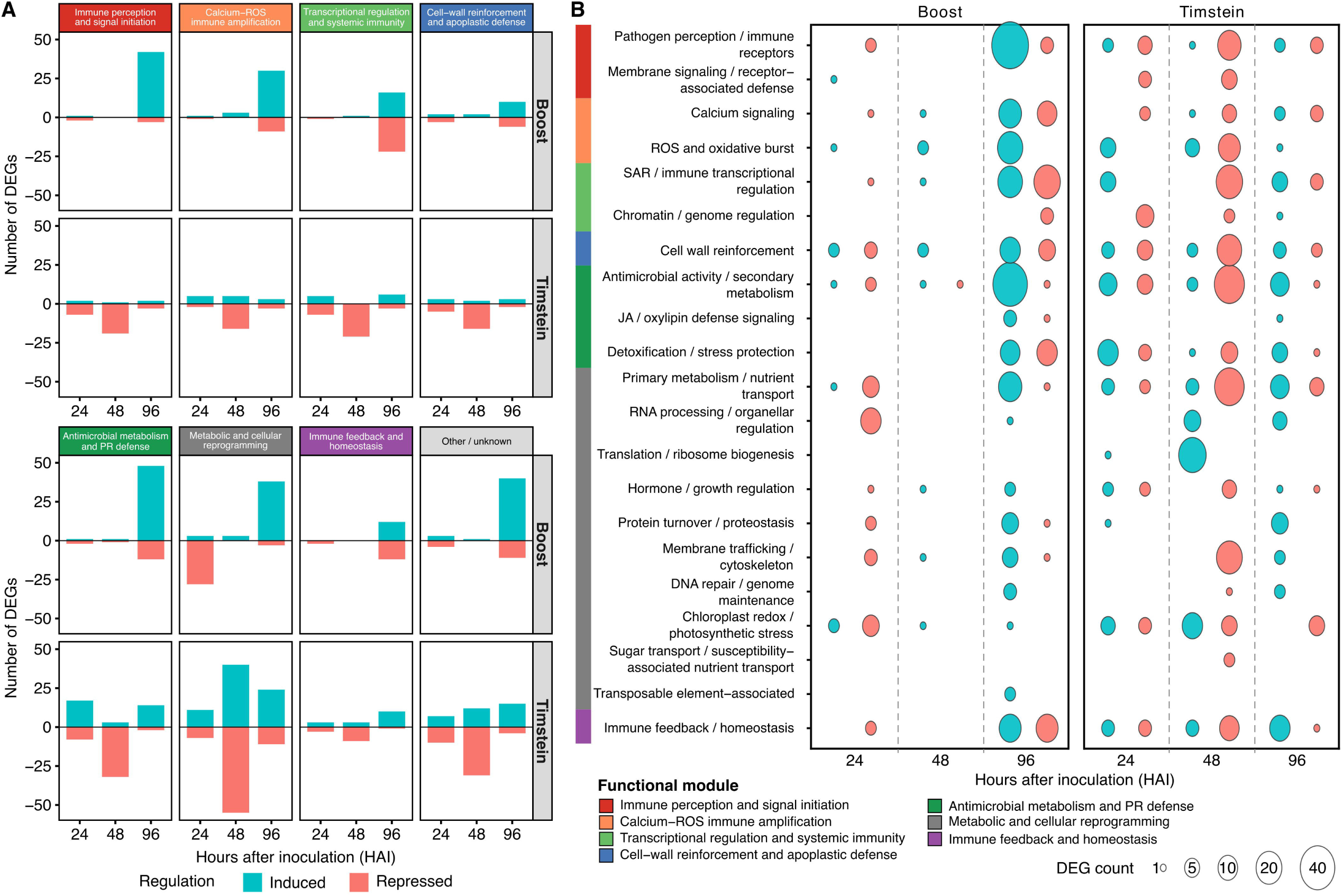
Contrasting temporal organization of immune processes in Boost and Timstein during bacterial leaf streak infection. **(A)** Numbers of induced and repressed DEGs assigned to each functional module in Boost and Timstein at 24, 48, and 96 HAI. Positive bars represent induced genes, and negative bars represent repressed genes. Modules include immune perception and signal initiation, calcium–ROS immune amplification, transcriptional regulation and systemic immunity, cell-wall reinforcement and apoplastic defense, antimicrobial metabolism and pathogenesis-related defense, metabolic and cellular reprogramming, immune feedback and homeostasis, and Other/unknown. **(B)** Comparative bubble plot showing the temporal regulation of detailed biological processes in Boost and Timstein. Bubble size represents the number of DEGs assigned to each process, and color represents the direction of regulation. Dashed vertical lines separate the 24, 48, and 96 HAI responses. The same functional classification framework was applied to both cultivars.

We integrated these comparative genomic and transcriptomic observations into the conceptual model (Figure 6). The model illustrates two contrasting immune trajectories following *Xtu* infection. In Boost, the receptor kinase-rich *QBls.fcu-3B.1* haplotype is associated with successful progression from pathogen perception to signal amplification and coordinated defense deployment. In Timstein, broad suppression of defense-related pathways during the critical 48 HAI transition is associated with disruption of this progression, resulting in a later but less coordinated transcriptional response. This model summarizes the contrasting immune trajectories identified in this study and provides a putative framework for future *in planta* functional validation of candidate genes within the *QBls.fcu-3B.1* interval for BLS response.

**Figure 6:**
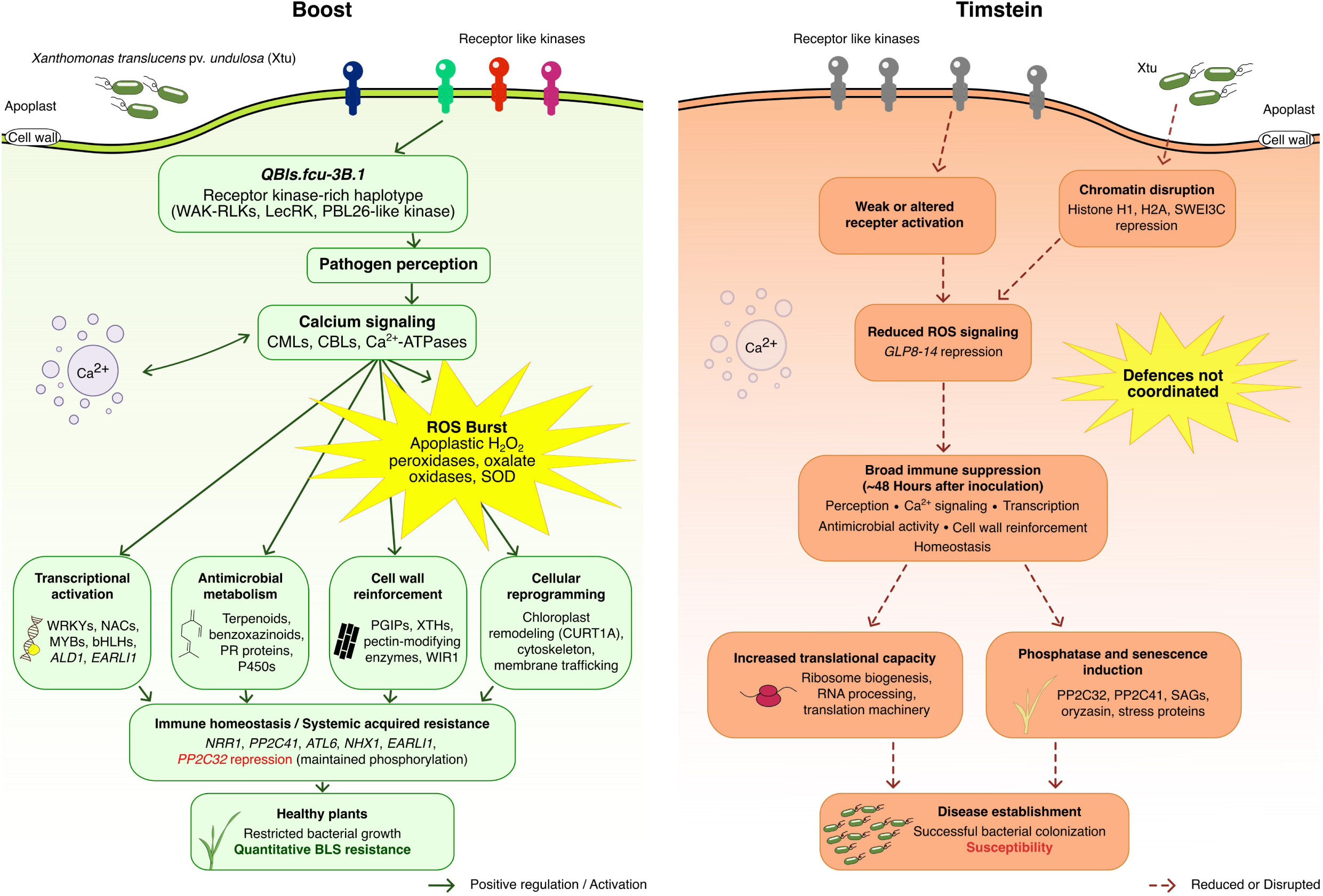
Proposed model for contrasting responses against BLS in Boost and Timstein. Working model integrating comparative genomic and time-resolved transcriptomic evidence for the resistant cultivar Boost and the susceptible cultivar Timstein following infection with *Xanthomonas translucens* pv. *undulosa*. In Boost, the receptor kinase-rich *QBls.fcu-3B.1* haplotype might contribute to pathogen perception. Receptor activation is followed by calcium signaling and ROS production, leading to transcriptional activation, antimicrobial metabolism, cell-wall reinforcement, cellular reprogramming, systemic acquired resistance, and immune homeostasis. These coordinated processes are associated with restricted bacterial growth and quantitative BLS resistance. In Timstein, altered receptor activation, reduced ROS-associated signaling, and repression of chromatin-associated genes preceded broad suppression of immune processes at approximately 48 hours after inoculation (HAI). Increased expression of translational machinery, phosphatases, and senescence-associated genes accompanies successful bacterial colonization and susceptibility. Solid green arrows indicate proposed activation or positive regulation, whereas dashed red arrows indicate reduced or disrupted processes. The model represents an evidence-supported framework and does not imply functional validation of individual genes or pathways.

## Discussion

For decades, efforts to understand disease resistance have been focused primarily on identifying R genes and structural variations within immune receptor families. This approach has been particularly successful for qualitative resistance (Nelson et al., 2018). However, most disease resistance in crop species is quantitative (Poland et al., 2009), arising from the combined action of numerous loci and complex regulatory interactions (Roux et al., 2014; Corwin and Kliebenstein, 2017). These perspectives suggest that receptor content alone may be insufficient to explain quantitative disease resistance, which involves genes acting across pathogen perception, signal transduction, hormone homeostasis, and metabolism (Gou et al., 2023). Our findings are consistent with this perspective. Despite possessing similar immune receptor repertoires, Boost and Timstein followed distinct transcriptional trajectories after BLS infection. Resistance in Boost might be associated with constitutive receptor expression and progression through an early signaling transition that preceded coordinated genome-wide defense activation.

Several candidate receptor kinases exhibited constitutively higher expression in the resistant cultivar Boost under both mock and BLS inoculated conditions, which suggests that the two haplotypes differ in baseline receptor expression rather than in infection-induced activation. A similar relationship between receptor kinase abundance and disease resistance has been reported for rice bacterial blight resistance gene *Xa3*/*Xa26*, where receptor expression in different genetic backgrounds determines both the level and spectrum of the resistance against *X. oryzae* pv. *oryzae* (Cao et al., 2007). In another study, heterologous expression of the *Arabidopsis* PRR EFR conferred broad spectrum bacterial resistance in both *Nicotiana benthamiana* and tomato without modification of downstream immune signaling (Lacombe et al., 2010). Although *QBls.fcu-3B.1* is a naturally occurring QTL rather than a transgene, our findings are consistent with the principle that enhanced receptor abundance increases immune surveillance before infection and influences subsequent immune responses. The evolutionary architecture of the candidate receptor kinases further supports this interpretation. Protein comparisons between syntenic homologous receptor kinases from Boost and Timstein revealed highly conserved intracellular kinase domains, whereas substantially greater sequence divergence elsewhere. This is consistent with the modular evolution of PRRs in which ligand-binding domains diversify under selection and intracellular signaling domains remain constrained to preserve signaling competence (Couto and Zipfel, 2016; Ngou et al., 2021). Our observations suggest that naturally occurring structural and regulatory variation within this receptor-rich haplotype may enhance pre-infection immune surveillance while avoiding the metabolic costs associated with constitutive defense activation (Huot et al., 2014).

Quantitative resistance appears to depend on successful progression through a critical intermediate stage of the immune response rather than on the magnitude of transcriptional change at any individual time point. Boost exhibited only 12 DEGs at 48 HAI (hours after inoculation), but these genes were enriched for calcium signaling, ROS production, transcriptional regulation, antimicrobial activity and early cell wall remodeling. These processes occupy central positions within the plant immune network that links pathogen perception to extensive downstream transcriptional reprogramming (Jones and Dangl, 2006; Marcec et al., 2019; Bhar et al., 2023). By 96 HAI, this focused signaling state was followed by coordinated activation of seven defense-related modules, consistent with progression from early signal amplification to broad immune deployment. In contrast, Timstein entered a transcriptionally suppressive state at 48 HAI, characterized by coordinated repression across defense-related modules. The contrasting trajectories observed at 48 HAI suggest that this intermediate transition represents a key stage during which resistant and susceptible outcomes become biologically distinct.

The susceptible interaction provides an equally informative perspective. The widespread suppression of functionally distinct defense processes reflects global failure rather than failure of a single immune pathway. The coordinate repression observed across immune perception, calcium-ROS signaling, transcriptional regulation, antimicrobial metabolism, and cell wall reinforcement suggests higher order regulatory reprogramming of the host responses. One of the most striking features of the susceptible interaction was the early repression of linker histone H1 genes preceding the extensive transcriptional suppression observed later during infection. Previous studies have indicated that bacterial pathogens manipulate host chromatin architecture and histone dynamics to reshape defense-associated transcription (Ma et al., 2011). Our data does not establish whether the chromatin-associated changes observed in Timstein result directly from effector activity, cultivar-specific regulatory differences or secondary consequences of infection. However, their early occurrence raises the possibility that alterations of chromatin organization contribute to establishment of the susceptible transcriptional state, which contrasts with a recent study reporting increased defense in plants with a mutant linker histone H1 gene (Sheikh et al., 2023). Further, in our study, the genes associated with ribosome biogenesis, RNA processing, and translation were simultaneously induced during the same infection timeframe. Pathogens are known to manipulate host translation processes during compatible interactions (Lee et al., 2023), although the functional significance of the transcriptional changes observed here remains to be determined. Taken together, these observations suggest that susceptibility is accompanied by extensive reorganization of host regulatory and cellular processes rather than simply a passive failure to activate defense.

Our findings suggest that quantitative BLS resistance may depend on pre-existing immune preparedness together with the capacity to sustain coordinated defense during infection. The natural variation at *QBls.fcu-3B.1* influences the baseline immune state of the host and affects the efficiency with which early pathogen perception progresses into coordinated downstream defense. In the susceptible interaction, failure to maintain this progression coincided with early chromatin-associated changes and subsequent broad transcriptional reprogramming, highlighting the importance of regulatory context in determining disease outcome. Together, these findings link natural genomic variation to immune dynamics and provide a putative framework for *in planta* functional validation and characterization of receptor-rich resistance loci and for genomics-assisted improvement of durable bacterial disease resistance in wheat.

## Methods

### Plant material and bacterial inoculation

The spring wheat cultivars Boost, which provides quantitative resistance to BLS, and Timstein, which is susceptible, were used in this study. We used a highly virulent *X. translucens* pv. *undulosa* strain P3 from North Dakota (Sapkota et al., 2018) for the bacterial infection. The bacterial cells were grown overnight washed from the plates and resuspended in 1× PBS buffer (300 ml) to an optical density of 0.6 (OD_600_) with a final pH of 7.4. The bacterial suspension was added with 5-6 drops of Tween20 and inoculated onto the plants using spray method (Acharya et al., 2024). The mock inoculum contained 300ml 1× PBS buffer with 5-6 drops of Tween20.

### DNA extraction, genome sequencing, and assembly of Boost

For DNA extraction, the plants were grown in a hydroponic system with controlled growth conditions (12 hours (h) of light). The temperature settings for day and night were set at 20 °C and 18 °C, respectively. The MaxiBloom® Hydroponics Plant Food from General Hydroponics, Sevastopol, CA, USA at 1.7 g/L was used to grow the plants. We collected the leaf samples for high molecular weight (HMW) DNA from 3-week-old plants. The HMW DNA was extracted using CTAB-Qiagen Genomic-tip method (Vaillancourt and Buell, 2019). After assessing the quality and quantity of DNA with a NanoDrop spectrophotometer and Qubit dsDNA HS assay, respectively, the HMW DNA was sheared to 17 kb fragments using Diagenode Megaruptor. The SMRTbell Express Template Prep Kit 2.0 and Sage BluePippin size selection kit (≥10Kb) were used to prepare sequencing libraries. PacBio HiFi sequencing was performed on the PacBio Revio platform using eight 30 h SMRT Cells with Sequel II Sequencing Kit 2.0, Sequencing Primer v5, and Sequel Binding Kit 2.2. We generated approximately 27× PacBio HiFi genome coverage data and assembled it into a primary contig assembly using hifiasm (v0.24) (Cheng et al., 2022) with default parameters for an inbred species (-l0). For scaffolding, chromosome conformation capture (Hi-C) libraries were produced from the same plant tissue used for PacBio sequencing and sequenced to generate Illumina paired-end reads at Phase Genomics (Seattle, WA). Using this Hi-C data, we scaffolded the primary assembly into chromosome-scale pseudomolecules using YaHS (v1.2a.2) (Zhou et al., 2023), followed by manual correction using JuiceBox (v2.15) (Durand et al., 2016). The IWGSC CS RefSeq v2.1 reference genome (The International Wheat Genome Sequencing Consortium (IWGSC) et al., 2018) was used to name and orient the pseudomolecules. BlobTools (v1.1.1) (Laetsch and Blaxter, 2017) was used to separate the organellar contigs and identify any other genomic contaminants. Further, we performed a telomere search on all the chromosomes using tidk (v0.2.31) (Brown et al., 2025) with the “clade” option set to “Poales” to search for “AAACCCT” telomeric repeats. The genome assembly was evaluated for accuracy, completeness, and contiguity, using tools such as QUAST (v5.2.0) (Gurevich et al., 2013), Merqury (Rhie et al., 2020), BUSCO (v 6.0.0) (Tegenfeldt et al., 2025), and LTR_retriever (v3.0.4) (Ou and Jiang, 2018).

### RNA extraction and sequencing

We performed the RNA-seq experiment on Boost and Timstein under control and BLS-treated conditions. Plants at the 2-to 3-leaf stage were spray inoculated with P3 strain of *Xtu* and upon inoculation. The plants were kept in the misting chamber for 48 h. The leaf tissue of 5-6 cm in length was collected at three time points, *viz.*, 24 hours after inoculation (HAI), 48 HAI, and 96 HAI with three biological replications, which resulted in a total of 36 samples for RNA sequencing. RNA extraction, library preparation, and sequencing was completed by Novogene Inc. (USA). The 150 bp paired end reads produced for all samples were checked for the strandedness using infer_experiment.py (v5.0.2) (https://rseqc.sourceforge.net/#usage-information) (Wang et al., 2012b). The RNA-seq data were unstranded and the downstream data analysis was performed accordingly.

### Repeat and Gene model annotations of Boost and Timstein

The transposable elements (TEs) were annotated with a well curated ClariTeRep database (Daron et al., 2014) following the parameters used in previous publications (Singh et al., 2024; Singh et al., 2026). The soft masking was performed with RepeatMasker (v4.1.5). We used RNA-seq and protein homology evidences for structural gene annotations using BRAKER3 (Gabriel et al., 2024) in ETP mode. The predicted protein coding genes were functionally annotated with InterProScan (v5.38-76.0) and putative gene functions were assigned based on domain conservation and sequence similarity with the curated UniProt-Swiss-Prot protein database (https://www.uniprot.org/help/downloads). The high-confidence (HC) and low-confidence gene classification was done following our previously published protocol (Singh et al., 2025). Further gene model completeness was assessed with BUSCO (v6.0.0) (Tegenfeldt et al., 2025). The naming convention for Boost (v1) and Timstein (v2) gene models was as follows: “*TraesBO1A01G*#######” and “*TraesTS1A02G*#######”, respectively. We used RGAugury (Li et al., 2016) (https://bitbucket.org/yaanlpc/rgaugury/src/master/) to predict resistance gene analogs in both Boost and Timstein genome assemblies with default parameters and databases such as pfam and InterProScan.

### Comparative genomics and synteny analysis

We compared the whole genome assembly of Boost with CS-IAAS_v1.1 (Liu et al., 2025) and Timstein (Singh et al., 2026) using chromeister (v1.5.a). The HC gene models from Boost and Timstein were used for synteny analysis. The all-vs-all protein alignments were generated with blastp and the synteny was computed with MCScanX (Wang et al., 2012a). The links were visualized using SynVisio (Bandi et al., 2022) with best reciprocal hits. The whole genome alignments were performed using minimap2 (v2.30-r1287) (Li, 2018). The structural variants (Supplementary Table S11 and S12) were identified between Boost vs CS-IAAS and between Boost vs Timstein using SyRI (v1.7.1) (Goel et al., 2019).

### Transcriptomics data analysis

We used fastp (v0.23.4) to trim reads and to assess the read quality (Chen, 2023). To minimize reference bias arising from structural divergence between cultivars, a dual reference mapping strategy was employed. The RNA-seq reads from Boost were aligned exclusively to Boost genome assembly, whereas Timstein reads were aligned to Timstein genome assembly using STAR (v2.7.6a) aligner (Dobin et al., 2013). Gene-level read counts were quantified using featureCounts and genes with fewer than ten reads in at least three samples were removed prior to downstream analysis. The differential gene expression analysis was performed separately for each cultivar using DESeq2 (v1.44.0) (Love et al., 2014) in R (v4.4.0). Principal component analysis, sample-to-sample distance clustering, and dispersion estimation were used to assess the data quality. Pairwise comparisons between BLS inoculated and mock samples were performed at each timepoint i.e., 24, 48, and 96 HAI. The genes with adjusted *P*-value < 0.01, absolute log_2_ fold change > 1, and mean normalized read counts ≥ 50 were considered differentially expressed genes (DEGs). The differential expression significance was assessed using the Wald test implemented in DESeq2 with the Benjamini-Hochberg correction for multiple testing. The figures were generated using ggplot2 (Wickham, 2016) in R.

### Gene Ontology (GO) enrichment and functional immune module classification

The GO annotations were extracted from InterProScan functional annotation results. The GO enrichment analysis of DEGs was performed using clusterProfiler (v4.12.2) (Yu et al., 2012). Enriched GO terms were identified using adjusted *P*-value < 0.05. The orgDB for Boost and Timstein are provided at https://doi.org/10.6084/m9.figshare.33201099.

To facilitate biological interpretation and direct comparison between resistant and susceptible cultivars, the DEGs were further organized in biologically curated immune response modules rather than solely relying on GO enrichment. The functional annotations were parsed using a hierarchical keyword-based classification pipeline implemented in R (https://doi.org/10.5281/zenodo.23027356), which assigns genes to detailed biological categories that were subsequently collapsed into seven immune modules representing successive stages of plant defense. The seven modules are: Immune perception and signal initiation, Calcium–ROS immune amplification, Transcriptional regulation and systemic immunity, Cell-wall reinforcement and apoplastic defense, Antimicrobial metabolism and PR defense, Metabolic and cellular reprogramming, Immune feedback and homeostasis. Each gene was assigned to a single highest-priority immune module to avoid redundancy. We applied the same framework to both cultivars to enable direct comparison of immune response dynamics.

### Candidate gene prioritization in QTL region and protein sequence analysis

Candidate genes within the major BLS resistance locus reported previously by Acharya et al. (2024) were prioritized. We used four criteria for prioritization: (i) physical location within the marker defined QTL interval, (ii) cultivar specific transcriptional response, (iii) immune receptor or defense related genes, and (iv) protein sequence divergence between Boost and Timstein. Orthologous proteins were identified using blastp searches and OrthoFinder (v2.5.5) (Emms and Kelly, 2019). Signal peptides, transmembrane domains, and conserved protein domains were predicted using SignalP (Teufel et al., 2022), DeepTMHMM (Hallgren et al., 2022), and InterProScan, respectively. Pairwise global protein alignments between Boost candidates and Timstein orthologues were generated using EMBOSS Needle (https://www.ebi.ac.uk/jdispatcher/psa/emboss_needle) and amino acid identity was calculated for complete proteins and individual functional domains to evaluate patterns of sequence conservation and divergence. Normalized expression values for prioritized candidate genes were extracted from DESeq2 for each cultivar. Mean normalized counts and standard error across biological replicates were calculated in R and visualized using ggplot2. All the final figures were assembled in Inkscape (https://inkscape.org).

## Data availability

The raw PacBio HiFi, Hi-C, and RNA-seq data have been deposited in NCBI under BioProject accession PRJNA1440611. The raw RNA-seq data are available in the NCBI Sequence Read Archive (SRA) under accessions SRR39754281-SRR39754316. The raw HiFi and Hi-C data are available in SRA under accessions SRR39753940-SRR39753950. The chromosome-scale Boost genome assembly has been deposited at DDBJ/ENA/GenBank under accession JCEPNZ000000000, and the version described in this manuscript is JCEPNZ010000000. The reference genome assembly is accessible through GenBank under GCA_062045534.1. The gene annotations, coding sequences, protein sequences, and high-and low-confidence gene models for Boost and Timstein (v2) are available through Figshare at https://doi.org/10.6084/m9.figshare.33201099. The Boost genome has also been deployed on GrainGenes genome browser with BLAST and genome browser functionality.

## Author contributions

R.G. and J.D.F. conceived the study. J.S., S.G., and R.G. designed the study. P.J.M. performed the genome assembly. J.S. performed genome annotation, comparative genomics, structural variant analysis, resistance gene analog annotation, and transcriptomic analyses, integrated the datasets, and prepared the figures. J.S. and S.G. wrote the original draft of the manuscript. S.G. contributed to experimental work, data generation, and manuscript revision. J.D.F., S.S., K.L.D.R., U.G., and Z.L. contributed to experimental design, resources, data interpretation, and manuscript revision. R.G. supervised the project, secured funding, and revised the manuscript. All authors read and approved the final manuscript.

## Acknowledgments

This research was supported by USDA Agricultural Research Service CRIS project number 7735-77100-046-000D. We thank the computational assistance provided by the Center for Computationally Assisted Science and Technology (CCAST) at NDSU and express gratitude to the Jack Dalrymple Agricultural Research Complex at NDSU for providing greenhouse and growth chamber facilities. Mention of trade names or commercial products in this publication is solely for the purpose of providing specific information and does not imply recommendation or endorsement by the U.S. Department of Agriculture. USDA is an equal opportunity provider and employer.

## Supplemental information (SI) Supplemental note

**Note S1: Chromosome-scale assembly and annotation of the Boost genome**

**Note S2: Additional structural features revealed by comparative analysis**

## Supplemental Figures

**Figure S1: Hi-C contact map of the chromosome-scale Boost genome assembly.** Heatmap showing Hi-C interaction frequencies across the 21 assembled wheat chromosomes. Strong interaction signals along the diagonal and within individual chromosomes indicate successful chromosome-scale scaffolding and correct ordering and orientation of contigs.

**Figure S2: BUSCO assessment of gene annotation completeness for Boost and Timstein.** BUSCO completeness for all predicted proteins, the high-confidence (HC) and low-confidence (LC) gene sets, and HC genes from the A, B, and D sub-genomes in Boost and Timstein.

**Figure S3: Whole-genome collinearity and structural comparison of the Boost genome assembly. (A)** Dot plot showing chromosome-level collinearity between Boost and the CS-IAAS wheat reference genome. **(B)** Dot plot showing chromosome-level collinearity between Boost and Timstein. The largely continuous diagonal patterns indicate extensive chromosome collinearity between genomes. **(C)** Detailed visualization of the chromosome 7B/2B translocation identified during comparative genome analysis.

**Figure S4: Quality assessment of RNA-seq datasets. (A)** Principal component analysis (PCA) of all RNA-seq libraries showing separation of samples according to cultivar, treatment, and sampling time. **(B)** Mean-dispersion relationship estimated by DESeq2 after normalization. Black points represent gene-wise dispersion estimates, the red curve indicates the fitted dispersion model, and blue points represent the final dispersion estimates used for differential expression analysis. **(C)** Hierarchical clustering heatmap of normalized expression values demonstrating high reproducibility among biological replicates.

## Supplemental Tables

**Table S1:** Summary of raw PacBio and Hi-C data produced in this study.

**Table S2:** Primary assembly statistics of Boost genome.

**Table S3:** Summary statistics of Boost genome assembly.

**Table S4:** Chromosome length (bps) comparison of Boost, Timstein, CS-IAAS v1.1, and IWGSC RefSeq v2.1.

**Table S5:** The QV and error rate of Boost chromosomes computed by Merqury.

**Table S6:** Summary of telomeric repeats found on each sub-genome of Boost and Timstein chromosomes.

**Table S7:** Summary of transposable elements (TEs) in Boost assembly.

**Table S8:** Comparison of multiple genic features of chromosome assigned high-confidence (HC) gene models between Boost and Timstein gene models.

**Table S9:** Summary of resistance gene analogs predicted from HC gene models in Boost and Timstein genome assemblies.

**Table S10:** Chromosome-wise breakdown of RGAs found in Boost and Timstein genome assemblies.

**Table S11:** Structural variants identified in Boost genome compared with CS-IAAS and Timstein genomes.

**Table S12:** Summary of inversions found in Boost compared to CS-IAAS and Timstein in each chromosome.

**Table S13:** Physical positions and probe information for the genetic markers defining the *QBls.fcu-3B.1* interval in the Boost and Timstein genome assemblies.

**Table S14:** Annotated genes located within *QBls.fcu-3B.1* interval in the Boost genome assembly.

**Table S15:** Annotated genes located within *QBls.fcu-3B.1* interval in the Timstein genome assembly.

**Table S16:** Detailed information about code names used in transcriptomics study.

**Table S17:** List of differentially expressed genes in *QBls.fcu-3B.1* in Boost vs Timstein comparisons at 24, 48, and 96 HAI.

**Table S18:** Domain comparison of five candidate genes of Boost and their orthologs in Timstein.

**Table S19:** Mapping statistics of Boost RNAseq data to Boost genome.

**Table S20:** Differentially expressed genes identified in Boost following BLS inoculation relative to mock-treated plants at 24, 48, and 96 hours after inoculation (HAI). Genes were considered differentially expressed at padj < 0.01 and |log2FC| > 1.

**Table S21:** Gene Ontology (GO) enrichment analysis of upregulated and downregulated genes in Boost following BLS inoculation at 24, 48, and 96 hours after inoculation.

**Table S22:** Selected differentially expressed genes in Boost for plotting and their functional classification.

**Table S23:** Mapping statistics of Timstein RNAseq data to Timstein genome.

**Table S24:** Differentially expressed genes identified in Timstein following BLS inoculation relative to mock-treated plants at 24, 48, and 96 hours after inoculation (HAI). Genes were considered differentially expressed at padj < 0.01 and |log2FC| > 1.

**Table S25:** Gene Ontology (GO) enrichment analysis of differentially expressed genes in Timstein following bacterial leaf streak (BLS) inoculation at 24, 48, and 96 hours after inoculation (HAI).

**Table S26:** Selected differentially expressed genes in Timstein for plotting and their functional classification.

