## Supplementary material for "Divergent immune trajectories distinguish resistance and susceptibility to bacterial leaf streak in wheat": Singh_et_al_2026_Boost_Supplemental_material.docx

**Note S1. Chromosome-scale assembly and annotation of the Boost genome**

The chromosome-scale genome assembly of the bacterial leaf streak-resistant wheat cultivar Boost was generated using multiple PacBio HiFi sequencing runs followed by Hi-C scaffolding. A detailed summary of the sequencing datasets, including read number, read length distributions, sequencing quality, and library statistics, is provided in Table S1. Assembly of the HiFi reads produced a primary contig assembly comprising 4,164 contigs with a total assembled length of 14.61 Gb, a contig N50 of 45.83 Mb, and the largest contig spanning 336.48 Mb (Table S2). Hi-C scaffolding anchored the assembly into 21 chromosome-scale pseudomolecules, producing a final genome assembly of 14.58 Gb, of which 14.33 Gb was assigned to chromosomes (Table S3). The final assembly showed high continuity with a scaffold N50 of 715.84 Mb, L50 of 10, and only 927 gaps remaining within chromosome sequences. Assembly quality assessment demonstrated high completeness and sequence accuracy, with BUSCO completeness of 98.2%, 97.7%, and 99.6% for the A, B, and D sub-genomes, respectively, a consensus quality value (QV) of 67.37, k-mer completeness of 98.79%, and LAI values ranging from 13.69, 17.45, and 18.10 for A, B, and D sub-genomes (Table S3).

Chromosome lengths of the Boost assembly were highly comparable to those of Timstein, CS-IAAS v1.1, and IWGSC RefSeq v2.1, with chromosome 3B representing the largest chromosome (Table S4). Repeat annotation showed that transposable elements occupy 85.82% of the Boost genome (Table S7). Class I retrotransposons represented the predominant repeat fraction, with LTR retrotransposons, particularly Gypsy and Copia elements, accounting for most repetitive DNA, consistent with the genomic organization of bread wheat. Genome-wide resistance gene analogue (RGA) annotation identified comparable immune receptor complements in the two cultivars (Table S9). Boost contained modestly more receptor-like kinases (3,137 vs. 3,020) and receptor-like proteins (460 vs. 428) than Timstein, whereas most other RGA classes were similar. Chromosome-wise distributions of all RGA classes are summarized in Table S10, providing a genome-wide framework for investigating immune receptor organization beyond the *QBls.fcu-3B.1* resistance locus.

Complete chloroplast and mitochondrial genome assemblies were also recovered from the sequencing data and are included as part of the Boost genomic resource (https://doi.org/10.6084/m9.figshare.33201099).

**Note S2: Additional structural features revealed by comparative analysis**

Whole-genome comparisons identified several chromosome-specific features that were not directly related to the bacterial leaf streak resistance loci but provide additional context for the Boost genome. Boost chromosome 7A was approximately 70 Mb shorter than the corresponding chromosomes of Timstein and CS-IAAS. Alignment of Boost HiFi reads to the distal region of the Chinese Spring chromosome provided limited support for this terminal segment, suggesting substantial cultivar-specific sequence divergence or absence of part of the corresponding region rather than an unsupported assembly extension. Gene-level collinearity analysis also detected a limited number of inter-chromosomal collinear blocks in addition to the expected chromosome-to-chromosome synteny. These included a reverse-oriented block containing 62 collinear genes between Boost chromosome 7B and Timstein chromosome 2B (Figure S3C). Because the present analyses cannot distinguish structural rearrangements from duplicated homoeologous segments, these observations are reported as comparative genomic features without inferring their evolutionary origin.

Detailed inspection of the *QBls.fcu-3B.1* interval identified structural variation beyond the receptor kinase candidates discussed in the main text. Across the interval, SyRI identified 8,968 SNPs, 483 deletions, 455 insertions, 27 duplications, one inversion, 16 translocations, 30 highly divergent regions, and additional complex structural rearrangements including inverted duplications and inverted translocations. Among these variants, a single canonical inversion was detected within the interval. The inversion overlapped the high-confidence Timstein gene *TraesTS3B02G0021800*, annotated as a putative coatomer subunit beta′-3, extending across the 5′ portion of the gene. No corresponding high-confidence gene was annotated at the homologous Boost coordinates. The biological significance of this inversion remains unknown. In addition to the receptor kinase clusters highlighted in the main text, the two haplotypes differed in the organization of several tandem gene families, including germin-like proteins, expansins, resistance-associated proteins, and genes of unknown function, further illustrating the localized structural complexity of the resistance locus.

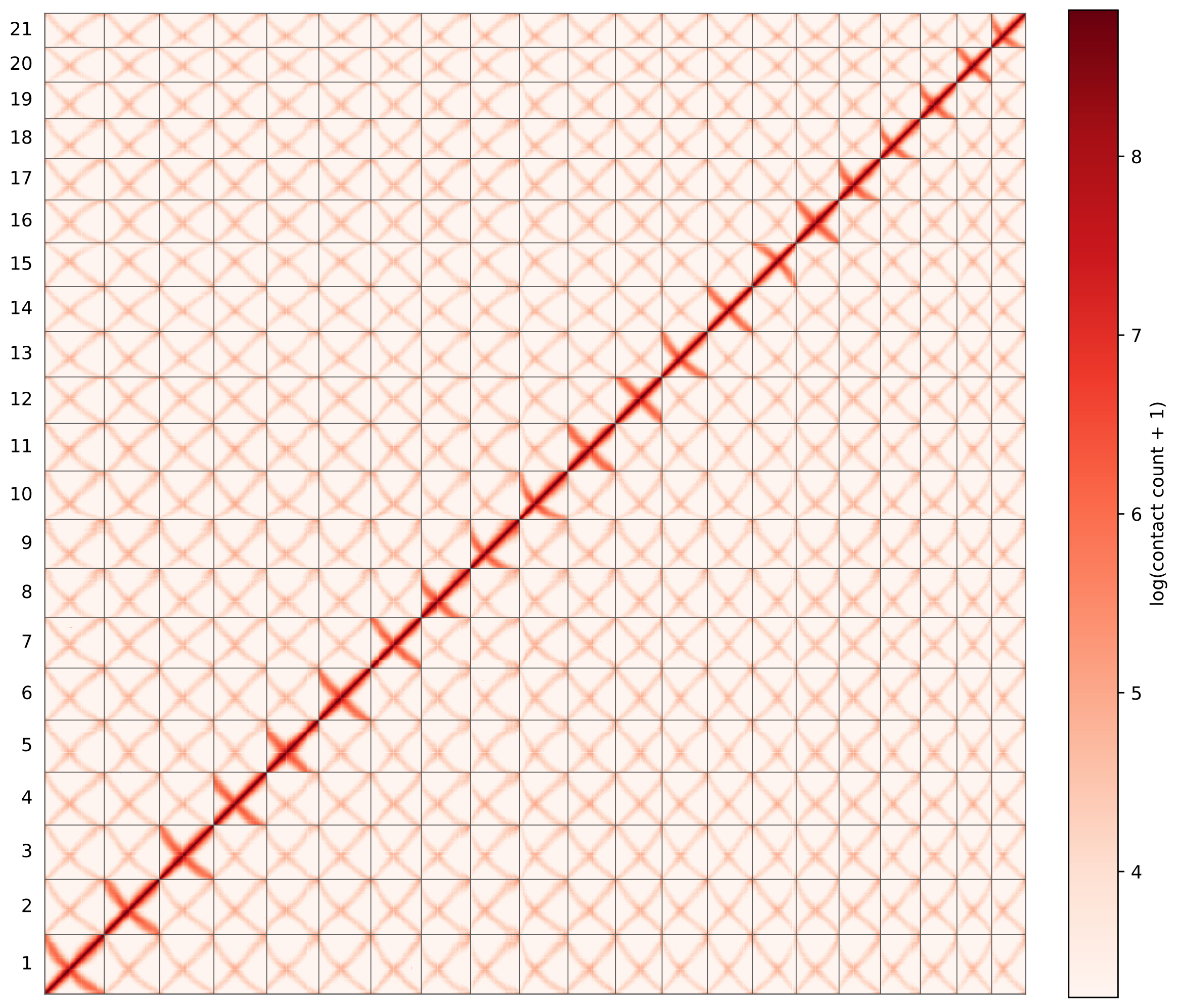

**Figure S1: Hi-C contact map of the chromosome-scale Boost genome assembly.** Heatmap showing Hi-C interaction frequencies across the 21 assembled wheat chromosomes. Strong interaction signals along the diagonal and within individual chromosomes indicate successful chromosome-scale scaffolding and correct ordering and orientation of contigs.

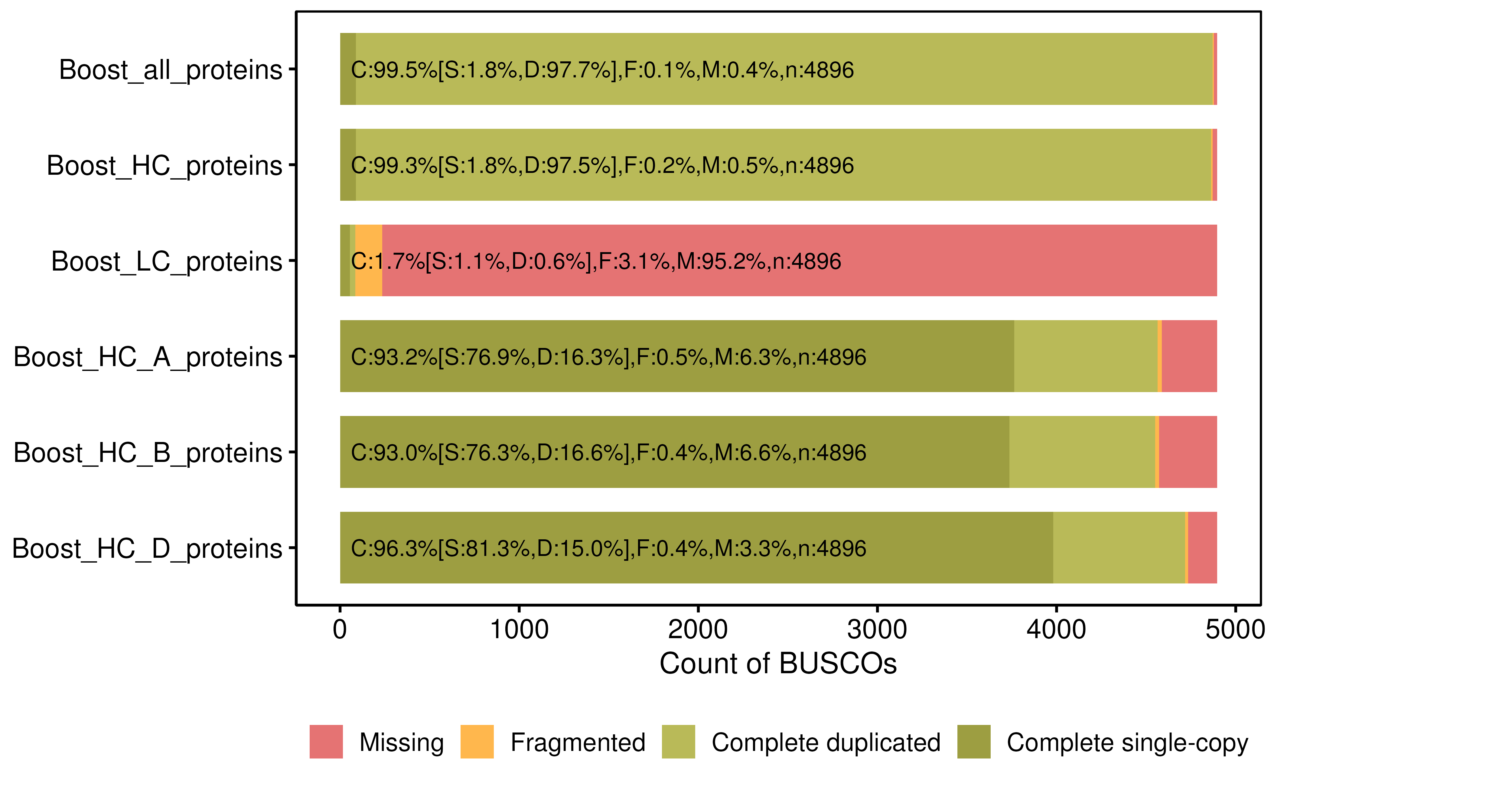

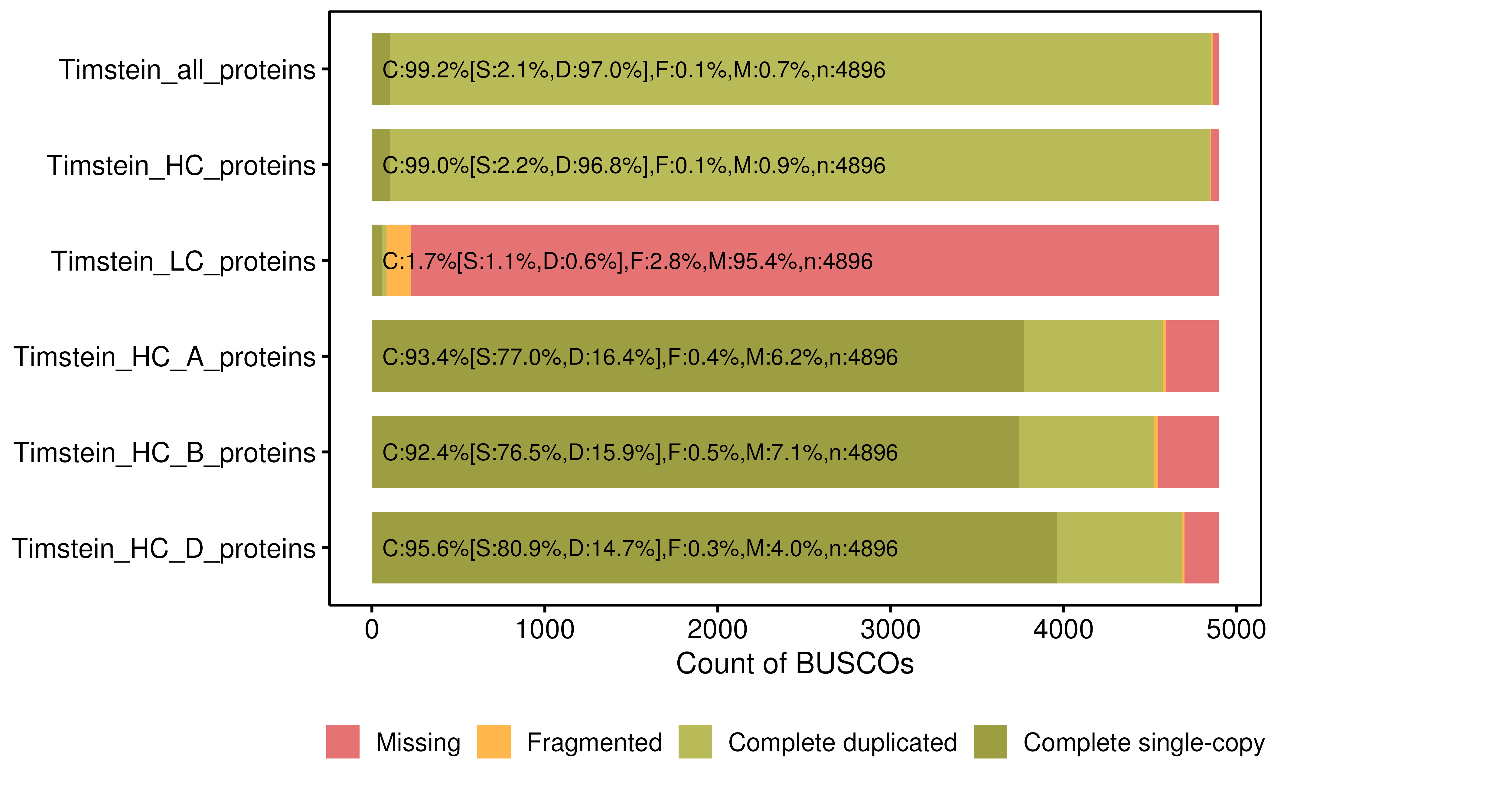

**Figure S2: BUSCO assessment of gene annotation completeness for Boost and Timstein.** BUSCO completeness for all predicted proteins, the high-confidence (HC) and low-confidence (LC) gene sets, and HC genes from the A, B, and D sub-genomes in Boost and Timstein.

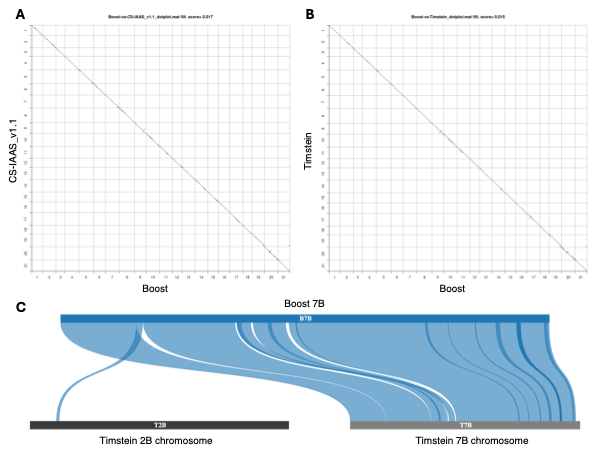

**Figure S3: Whole-genome collinearity and structural comparison of the Boost genome assembly. (A)** Dot plot showing chromosome-level collinearity between Boost and the CS-IAAS wheat reference genome. **(B)** Dot plot showing chromosome-level collinearity between Boost and Timstein. The largely continuous diagonal patterns indicate extensive chromosome collinearity between genomes. **(C)** Detailed visualization of the chromosome 7B/2B translocation identified during comparative genome analysis.

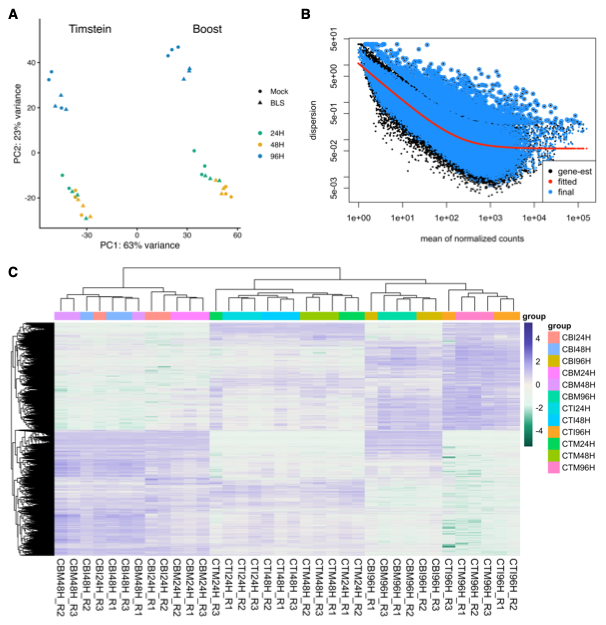

**Figure S4: Quality assessment of RNA-seq datasets. (A)** Principal component analysis (PCA) of all RNA-seq libraries showing separation of samples according to cultivar, treatment, and sampling time. **(B)** Mean-dispersion relationship estimated by DESeq2 after normalization. Black points represent gene-wise dispersion estimates, the red curve indicates the fitted dispersion model, and blue points represent the final dispersion estimates used for differential expression analysis. **(C)** Hierarchical clustering heatmap of normalized expression values demonstrating high reproducibility among biological replicates.
